# Novel cryo-electron tomography structure of Arp2/3 complex in cells reveals mechanisms of branch formation

**DOI:** 10.1101/2020.08.25.266262

**Authors:** Florian Fäßler, Georgi Dimchev, Victor-Valentin Hodirnau, William Wan, Florian KM Schur

## Abstract

The actin-related protein (Arp)2/3 complex nucleates branched actin filament networks pivotal for cell migration, endocytosis and pathogen infection. Its activation is tightly regulated and involves complex structural rearrangements and actin filament binding, which are yet to be understood. Here, we report a 9.0Å resolution structure of the actin filament Arp2/3 complex branch junction in cells using cryo-electron tomography and subtomogram averaging. This allows us to generate an accurate model of the active Arp2/3 complex in the branch junction and its interaction with actin filaments. Our structure indicates a central role for the ArpC3 subunit in stabilizing the active conformation and suggests that in the branch junction relocation of the ArpC5 N-terminus and the C-terminal tail of Arp3 is important to fix Arp2 and Arp3 in an actin dimer-like conformation. Notably, our model of the branch junction in cells significantly differs from the previous *in vitro* branch junction model.

## Introduction

Spontaneous actin filament polymerization is an energetically unfavourable process and nucleation factors are required to overcome this rate-limiting step in actin filament assembly. The heptameric 224 kDa Arp2/3 complex, consisting of the two actin-related proteins 2 and 3 (Arp2 and Arp3) and five additional subunits ArpC1-5, generates branched actin networks by inducing the formation of so-called actin filament Arp2/3 complex branch junctions on pre-existing actin filaments.

The Arp2/3 complex plays a role in both physiological and pathological processes, such as lamellipodial protrusion, vesicle trafficking and also the motility of various pathogens. Arp2/3 complex activation is tightly regulated, requiring multiple regulatory proteins, including nucleation promoting factors (NPFs), ATP and monomeric actin [1]. NPFs, such as members of the Wiskott-Aldrich syndrome protein (WASP) and SCAR/WAVE families, induce the structural changes required for Arp2/3 complex activation via their WASP homology 2, connector, and acidic (WCA) domains, where the CA motifs bind the Arp2/3 complex and the W motif delivers the first actin monomer [2].

The available structural and biochemical information of the Arp2/3 complex and its interactions with other proteins have been mostly obtained via a reductionist approach, i.e. studying the Arp2/3 complex *in vitro* in isolation, bound to NPFs or their fragments [3,4], or stabilizing and destabilizing proteins, including cortactin [3], coronin or Glia maturation factor (GMF) [5,6]. The conformation of the Arp2/3 complex within the branch and its interaction with the mother and daughter filament were largely derived from fitting high-resolution x-ray crystallography models of the inactive complex [7,8] into low-resolution *in vitro* electron microscopy (EM) and electron tomography (ET) reconstructions [9–11], as well as molecular dynamics (MD) simulations [12]. These studies led to the model that upon activation, the Arp2 and Arp3 subunits reposition into a short-pitch conformation resembling an actin filament-like state, providing the basis for further actin polymerization. In the *in vitro* branch junction model all seven subunits were reported to form contacts with the mother filament upon branch formation [9,11]. Binding of the Arp2/3 complex to the mother filament was also proposed to lead to a conformational change in the filament, increasing branch junction stability [9,12]. However, due to the low-resolutions at ^~^25Å of the previously determined branch junction models, the currently suggested molecular contacts of the active Arp2/3 complex in the branch junction and the associated changes in the mother filament remain ambiguous. Specifically, an accurate structure of the actin filament Arp2/3 complex branch junction has been lacking; such a structure would better describe the functional consequences of the exact interfaces in the branch junction on its formation and stability.

Cryo-electron tomography (cryo-ET) combined with image processing approaches such as subtomogram averaging (STA) can provide structural insights into native cellular environments [13]. Medium to high-resolution structures of proteins within their native environment using these methods have so far been only described for a few specimens, often large or highly symmetrical assemblies[14–18]. Among the limitations that restrict the resolution that can be obtained in cells are the thickness of the specimen, reduction of which often requires additional specimen preparation steps, and the number of protein complexes available for generating a higher resolution structure [13].

The lamellipodium is a thin sheet-like membrane extension at the leading edge of migrating cells, which has a height of 100-200 nm and is densely filled with an Arp2/3 complex-dependent branched actin filament network (also referred to as dendritic network). It has been a prominent model to study actin network topology and ultrastructure using both room-temperature and cryo-electron microscopy approaches [19–22]. Due to the large number of actin filament Arp2/3 complex branch junctions within lamellipodia, we considered them the optimal target to determine the structure of branch junctions in their native environment at the highest possible resolution and to provide the outstanding answers to the important aforementioned open points on branch junction formation and stability.

## Results

### Cryo-ET of actin filament Arp2/3 complex branch junctions in fibroblast lamellipodia

We performed cryo-ET of lamellipodia of NIH-3T3 fibroblast cells, which we transiently transfected with a constitutively active variant of the small GTPase Rac (L61Rac). Rac overexpression has been previously reported to enhance lamellipodia formation by activating NPFs [20,23], facilitating the selection of appropriate regions for cryo-ET data acquisition. In order to further optimize conditions for image processing we used established extraction and fixation protocols involving the actin-stabilizing toxin phalloidin, allowing for a better visualization of actin filaments and bound complexes [20,22]. In agreement with previous observations, extraction and fixation did not change the ultrastructure of the actin network, and branch junctions were clearly retained within lamellipodia (Fig. S1, Movie S1).

Using a combination of semi-automatic particle detection, classification, STA and multiparticle refinement [18] we obtained a 9.0Å resolution structure of the branch junction from aligning and averaging 14,296 subtomograms (Fig. 1a, Fig. S2 and Movie S2, see Table S1 and *Materials and Methods* for details). At this resolution all subunits of the Arp2/3 complex and also individual actin monomers of the mother (designated here as M1-M8) and daughter filaments (D1-3) are clearly resolved.

**Fig. 1:**
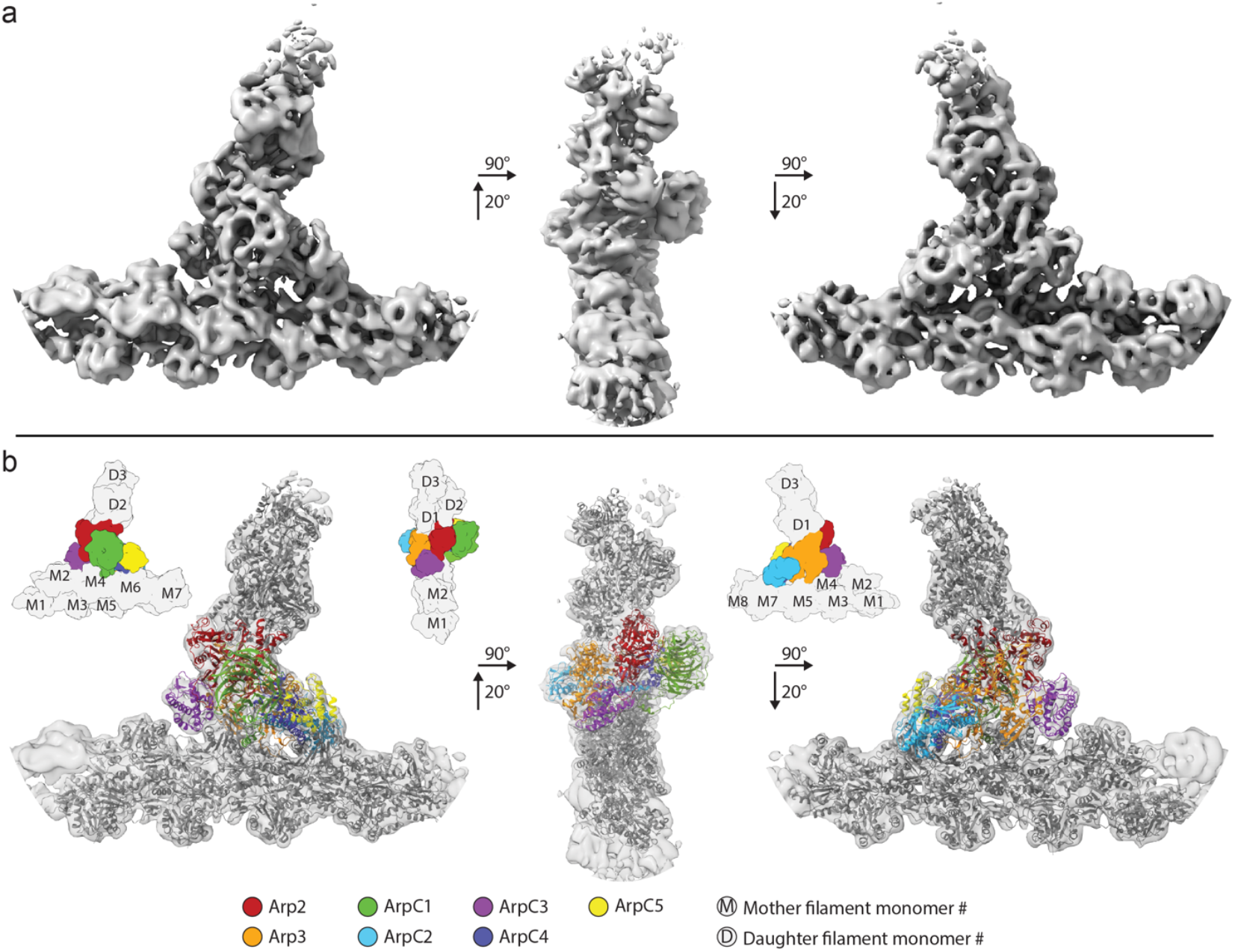
Subnanometer Structure of the actin filament Arp2/3 complex branch junction in cells. **(a)** Isosurface representation of the actin filament Arp2/3 complex branch junction in cells at 9Å resolution. The structure is shown from three orientations. A guide for orientation is given in (b). **(b)** The electron microscopy density map (shown transparent) with the flexibly fitted models of the Arp2/3 complex subunits and actin filaments. Schematic guides indicate the positions of the individual subunits of the complex and the monomers of both the mother and daughter filaments. A colour legend provides the color code for the fitted models. The color code and legend are used throughout the manuscript to aid the reader.

Moreover, visible secondary structure details corresponding to alpha-helices and various loops allowed an unambiguous placement of available high-resolution models of the individual Arp2/3 complex subunits and actin monomers into our density, without requiring any large-scale modifications of their tertiary structure. Our structure also allowed us to model the backbone trace for some regions in the Arp2/3 complex that were not resolved in crystal structures (Table S2). We then used MD-based fitting [24] to remove steric clashes and to generate a complete actin filament Arp2/3 complex branch junction model (Fig. 1b, Fig. S3, Movie S3 and Tables S2 and S3). The density corresponding to ArpC5 suggested increased flexibility at its N-terminus (Fig. S3). This flexibility could be due to alternate conformations of this region of the subunit, caused by the existence of the two isoforms ArpC5 and ArpC5L, in line with a recent study [25]. Hence, we restrained the movements of this subunit in the MD modelling. The quality of the reconstruction was further underlined by the presence of densities not occupied by either the Arp2/3 complex or actin, but at the exact location at which Phalloidin would be expected (Fig. S3). The fact that we were able to obtain subnanometer resolution in large part of our structure including the first actin monomers of the daughter filament indicates rigidity of the branch junction. We determined the angle of the branch junction in our structure to be 71 degrees, which is in agreement with some, but different to other previously reported *in vitro* and *in situ* studies, which showed branch junction angles ranging from 67 to ^~^80 degrees [9,11,12,20,26,27]. This led us to examine in more detail the structural changes occurring in the Arp2/3 complex upon branch junction formation and to compare our model to previously suggested conformations of the branch junction [9,12,28].

Specifically, we sought to understand the contribution of the different subunits to branch junction conformation and stability, in particular their contact with both the actin mother and daughter filament.

### The structure of the active Arp2/3 complex

First, we compared our active Arp2/3 complex model in the branch junction to a model of the soluble inactive complex obtained by x-ray crystallography [8] (Fig. 2). The biggest change upon branch junction formation was the actin-like heterodimer arrangement of Arp2 and Arp3, which resembles the first two subunits of the daughter filament (Fig. S4). In this regard our structure of the activated Arp2/3 complex in branch junctions agrees with a recently published *in vitro* structure of the NPF Dip1-activated *S. pombe* Arp2/3 complex that generates unbranched actin filaments [29] (Figure S4, Figure S5a,b). Also, in both structures of the activated Arp2/3 complex, ArpC2 and ArpC4 form a stable core and define the centre of rotation and translation of two subcomplexes consisting of ArpC2, Arp3, ArpC3 and ArpC4, ArpC5, ArpC1, Arp2 that move against each other to form the short pitch conformation (Movies S4 and S5).

**Fig. 2:**
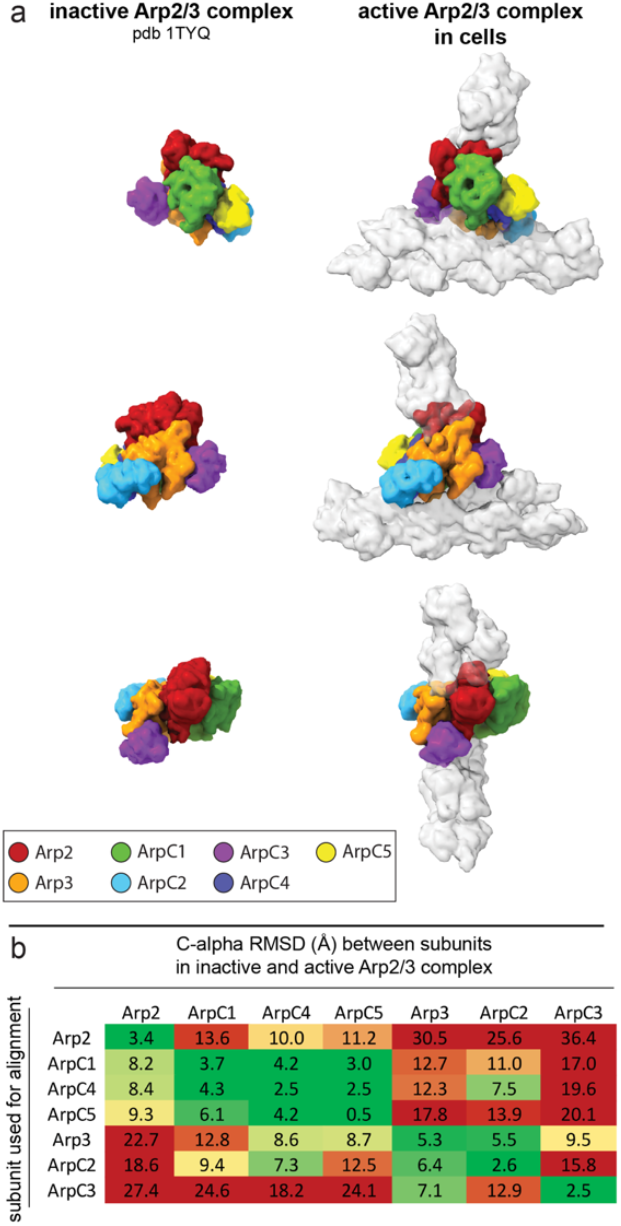
Comparison of Arp2/3 complex in its inactive conformation and in the branch junction in cells. **(a)** Molecular models of the Arp2/3 complex in the inactive (derived from pdb 1TYQ) and the active conformation shown as density maps filtered to 9.5Å resolution. The models are shown from three orientations, corresponding to the views in figure 1. **(b)** RMSD values (in Å) calculated between the inactive and active conformations of the individual Arp2/3 subunits (based on the models used in (a)). Rows indicate which subunit was used for aligning the full models against each other, prior to measurements between individual subunits of the inactive and active Arp2/3 complexes (indicated in the columns). The RMSD analysis reveals that the structural transition upon complex activation is accommodated by two subcomplexes consisting of Arp2, ArpC1, ArpC4 and ArpC5 and Arp3, ArpC2 and ArpC3, respectively, that rotate against each other along an axis formed by the large helices of ArpC2 and ArpC4 (Movie S4). RMSD variations between the same subunits in the inactive and active conformations derive from changes upon MD-based modeling of the x-ray crystal structure-derived model into the electron microscopy density map of the branch junction.

While in the case of unbranched filaments, nucleated by the Dip1-activated Arp2/3 complex, this structural change is induced by binding of Dip1 to ArpC4, other activating mechanisms must exist to induce this conformation in the branch junction, as the ArpC4 binding site is occupied by the mother filament (see below).

The structure of the activated Arp2/3 complex in the branch junction also reveals differences in inter-subunit contacts and distances compared to the Dip1-activated complex (Figure S5a,b). Specifically, in the branch junction we observe a significant movement of ArpC3 towards the D-loop of Arp2 providing additional stabilization to the active conformation (Fig. 2a,b, Fig. S5b).

### Actin filament Arp2/3 complex interactions

Fitting of a model of filamentous actin (pdb 6T20) [30] into our structure showed good correspondence, which however could be improved by refining the fit of the individual monomers, which led to small changes from their original filamentous conformation (Fig. S6, a and b). These changes however, did not affect the monomer conformation (1.76Å mean RMSD between C-alpha atoms of monomers in the mother filament and pdb 6T20) (Fig. S5c), in contrast to what has been predicted from an earlier low-resolution EM structure of the branch junction [9].

To derive a more accurate description of interaction interfaces between the Arp2/3 complex and the mother actin filament, we probed for C-alpha atoms of Arp2/3 complex subunits and actin monomers being in vicinity closer than 10Å (Fig. 3 and Table S4). In contrast to the currently accepted branch junction model derived from a low-resolution structure [9], only 5 subunits, namely Arp3, ArpC1, ArpC2, ArpC3 and ArpC4 contact the mother filament (Fig. 3a, Fig. S5a). ArpC5 does not form any contact with either the mother or the daughter filament, in line with previous reports suggesting its role to be mainly regulatory [25]. Correspondingly, in total 5 monomers (M2, M4-M7) in the mother actin filament are positioned to contact the Arp2/3 complex. A prominent interaction surface between the complex and the mother filament is formed by ArpC2 and ArpC4 contacting M5, M7 and M6, respectively (Fig. 3, a and b).

**Fig. 3:**
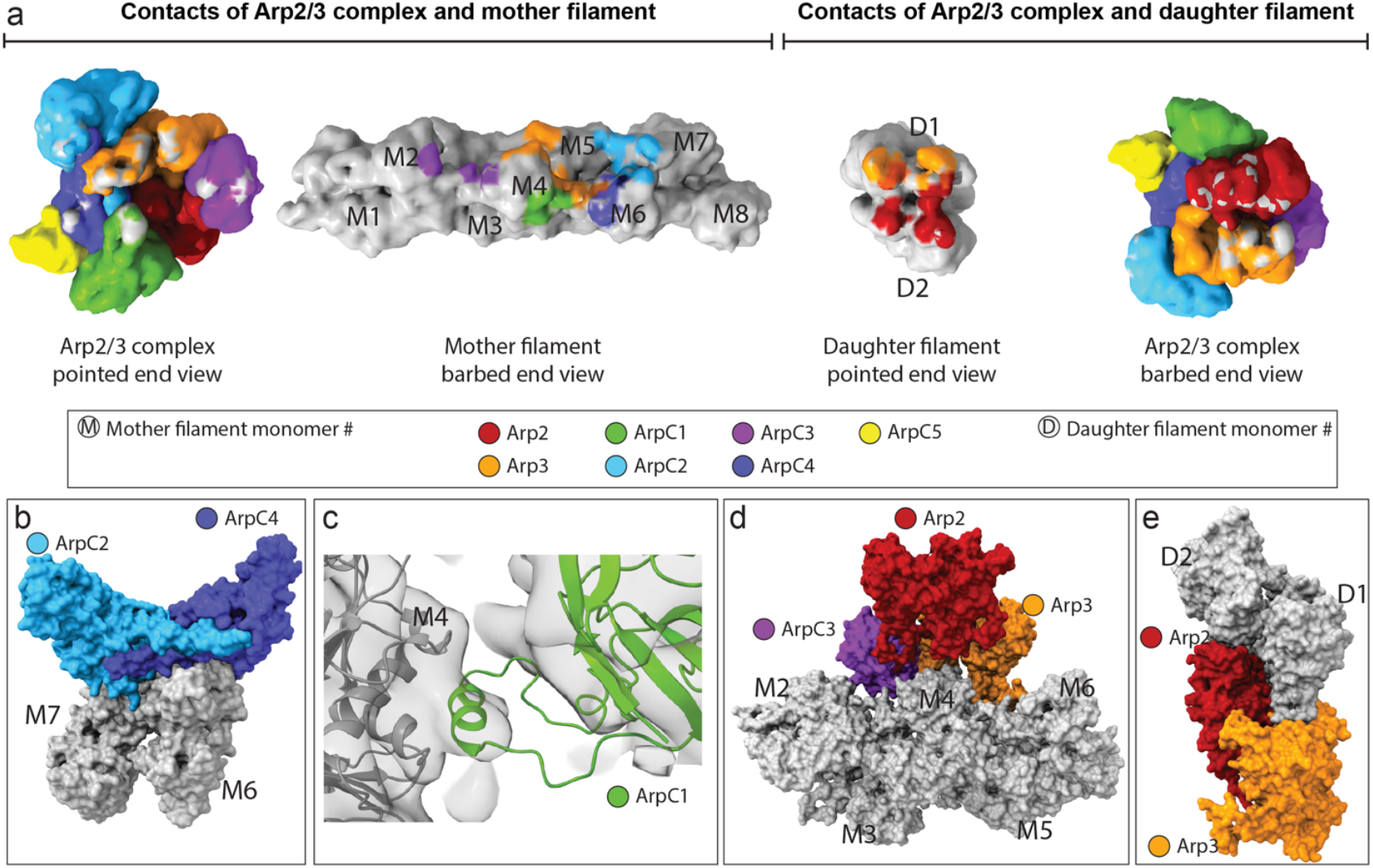
Actin-Arp2/3 complex interaction surfaces within the branch junction. **(a)** Interaction surfaces between the Arp2/3 complex and the actin mother and daughter filament. The surfaces for the Arp2/3 complex, mother and daughter filament are shown as density maps at 9.5Å resolution generated from their respective models. Grey coloring on the surface of Arp2/3 subunits indicates contact sites with actin and coloring of actin monomers in a specific color indicates a contact site with the associated Arp2/3 subunit (color code is given within the figure). Coloring was applied via the color zone command in ChimeraX in a 5.5Å radius around each C-alpha atom of the underlying model, which was positioned in a 10Å radius to a C-alpha atom of its putative interactor. **(b)** Surface representation of the interaction of ArpC2 and ArpC4 with the mother filament monomers M6 and M7 (the interaction of ArpC2 with M5 is omitted for clarity). **(c)** The protrusion helix of ArpC1 fitted into its density close to subdomains 1 and 3 of M4. **(d)** Surface representation of the interactions of Arp3 and ArpC3 with the mother filament. Note the cavity below Arp2, where no contacts between the Arp2 subunit and the mother filament are observed. ArpC3 acts as a linker between Arp2 and the mother filament. **(e)** Surface representation of the interaction between Arp2, Arp3 and the first two monomers of the daughter filament.

This is in agreement with biochemical evidence showing the importance of some of the residues involved in this interaction to nucleation and branch junction stability [28]. A density close to subdomains 1 and 3 of M4 fits to the previously proposed interaction of the ArpC1 protrusion helix (residues 297-305) with the mother filament (Fig. 3, a and c).

Arp3 establishes contacts with three mother filament monomers (M4-6), mostly via its D-loop and the base of subdomain 4 (Fig. 3, a and d). Additionally, ArpC3 does not only bind Arp2 and stabilizes the short pitch conformation, but also acts as a bridge between Arp2 and the mother filament, by forming small interfaces with actin monomers M2 color and M4 (Fig. 3, a and d). This central role of ArpC3 in the formation and maintenance of the branch is in accordance with previous studies showing that ArpC3 KO mice and *S. pombe* are not viable [31,32].

A cavity below Arp2 (Fig. 3d and Movie S3) highlights the overall loose contacts between the Arp2/3 complex and M4 and M5. This in line with our observation that formation and maintenance of a stable actin filament Arp2/3 complex branch junction do not require significant changes in actin monomers of the mother filament. This indicates that the mother filament does not need to adapt an unfavourable high-energy state to be primed for attachment of the Arp2/3 complex to its side, which might have implications for debranching or competition of actin regulatory factors with the Arp2/3 complex for the same binding site on the mother filament.

Arp2 and Arp3 are the only two subunits forming interfaces with the daughter filament (Fig. 3, a and e), confirming that the stability of connecting the activated Arp2/3 complex to the newly growing daughter filament is solely accomplished by actin-like interfaces [9,29].

### Arp2/3 complex activation and branch junction stabilization

Arp2/3 complex activation during branch junction formation requires the concerted action of two WCA binding events on two distinct sites on the Arp2/3 complex, specifically on Arp2-ArpC1 and Arp3 [33]. We asked if the described binding sites are accessible in our structure and if so, their location within the branch junction allows deriving further information about branch junction formation. A recent single-particle cryo-EM study of the inactive Arp2/3 complex bound to two WCA peptides showed that their C-helices interacted with structurally similar regions in both Arp2-ArpC1 and Arp3 [4]. Previously it was also suggested that for Arp2/3 complex activation, NPF binding to the Arp3 binding site requires the Arp3 C-terminal tail to be displaced from its position in the inactive conformation of the Arp2/3 complex, releasing its autoinhibitory function [4,34] and also eventually allowing the D-loop of the first daughter filament actin monomer to bind the Arp3 barbed end [29]. This agrees with our structure where no density corresponding to the C-terminal tail in its inactive conformation is present. Instead, our structure contains a density that places the Arp3 C-terminus in a position where it is flipped towards ArpC4 and Arp2 (Fig. 4a).

**Fig. 4:**
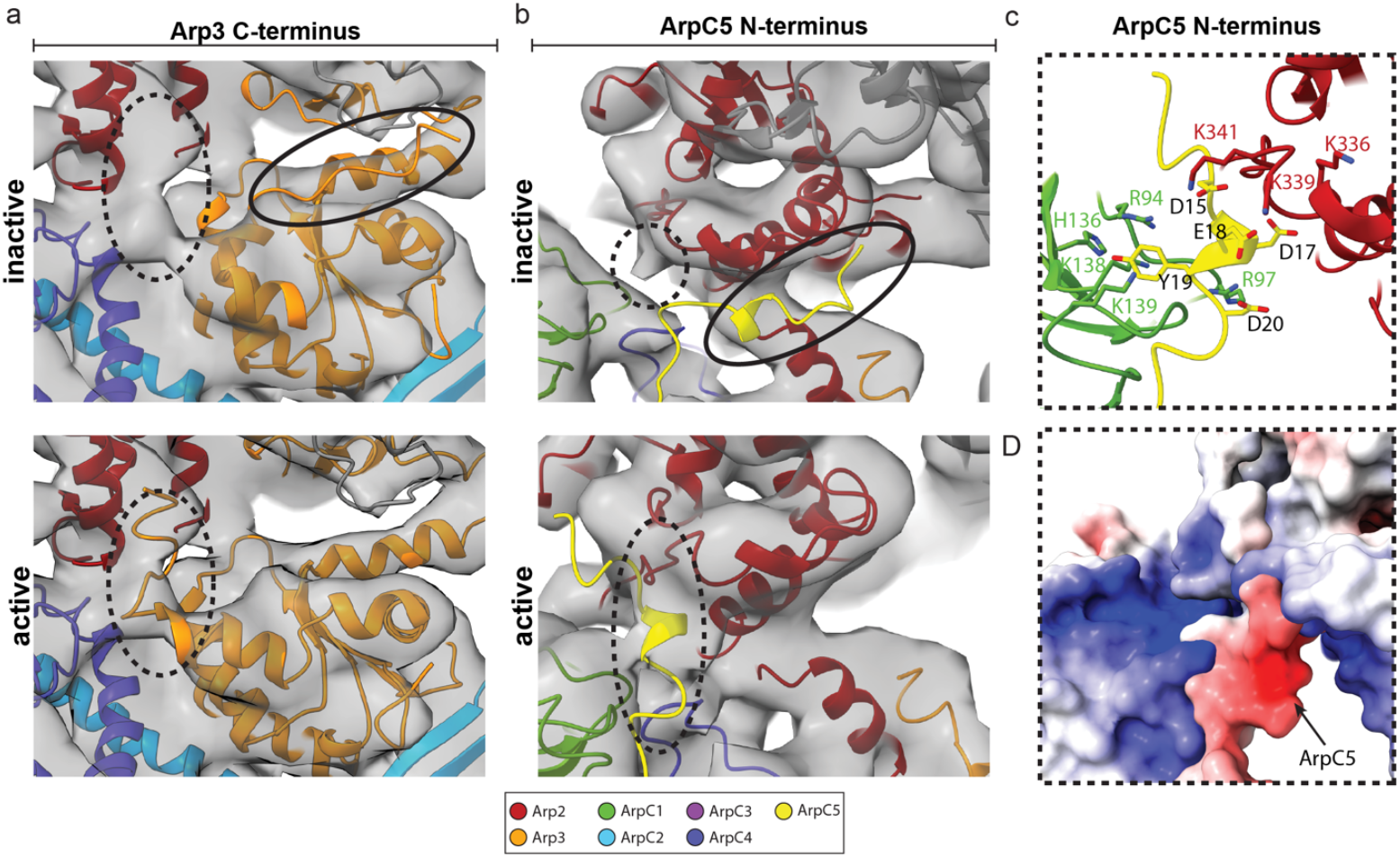
Structural changes in Arp3 and ArpC5 upon branch junction formation. **(a)** Comparison of the conformation of the Arp3 C-terminal tail in the inactive and active conformation. No density is observed for the C-terminal tail of Arp3 in its inactive conformation (top). Instead, the C-terminal tail can flip towards Arp2 and ArpC4, where it is accommodated by an empty density present in the branch junction structure (bottom). **(b)** No density is observed for the ArpC5 N-terminus at its binding side in the inactive complex (top). Instead, the ArpC5 N-terminus can be fitted into a density between ArpC1 and Arp2. **(c)** Positively charged residues in ArpC1 and Arp2 could coordinate the negatively charged N-terminus of ArpC5. **(d)** Electrostatic potential map of the area shown in (c).

In the inactive complex and the recent *in vitro* structure of the Dip1-activated *S. pombe* Arp2/3 complex, the ArpC5 N-terminus has been shown to bind the side of subdomain 3 of Arp2 [7,25,29]. In our structure, no density for this interaction can be observed (Figure 4b). It is therefore possible that upon branch junction formation, the ArpC5 N-terminus disengages from this binding site on the Arp2/3 complex. In our structure, faint densities were appearing between two strongly positively charged regions of Arp2 (involving residues K299, K336, K339, K341, R343) and ArpC1 (R94, R97, H136, K138, K139) some of which have been implicated in binding the D/E loop in the WCA motif of NPFs [4]. As NPFs dissociate from the Arp2/3 complex upon branch junction formation [10,35], we assumed it to be unlikely that these densities can be attributed to residual NPF branch junction binding.

Instead, the length of the linker connecting the ArpC5 N-terminus to its central helical core and the size of the empty density in our branch junction structure would allow the N-terminus to be fitted in this empty density between ArpC1 and Arp2 (Fig. 4b). Specifically, the negatively charged sequence of the N-terminus of ArpC5 (D15, D17, E18, Y19, D20) provide neutralization to the strongly positively charged region between ArpC1 and Arp2 (Fig. 4,c and d).

## Discussion

Taken together, these observations tempt us to propose the following model for stabilizing the branch junction and specifically the short pitch conformation. Actin-NPF binding to the high affinity binding site on Arp2-ArpC1 induces the short pitch conformation, which requires a contact between Arp2 and ArpC3 and also actin filament binding for stabilization. This could be further stabilized by the ArpC5 N-terminus substituting CA binding upon release of the NPF, which is required before nucleation can occur [36]. Recently, the mammalian Arp2/3 complex subunit isoforms ARPC1a, ARPC1b, ARPC5 and ARPC5L were shown to assemble into functionally different complexes in defined combinations, resulting in significantly changed actin network dynamics, partially caused by their varying interaction with the NPF cortactin and the branch junction severing factor coronin 1b [37]. The binding site of the ArpC5 N-terminus partially overlaps with the previously proposed binding site of cortactin [3]. The differential regulatory activity of ArpC5 isoforms could be caused by their varying binding strength of their N-terminus to the active conformation of the Arp2/3 complex, playing a regulatory role in interactions with cortactin, specifically selecting for Arp2/3 complexes of ArpC1/ArpC5 subunit isoform composition [38]. Additionally, activation of the Arp2/3 complex is associated with ATP binding, the binding of the C-helix of the second NPF to Arp3, and the release of the Arp3 C-terminal tail from its binding site in the inactive complex [4,29,34]. The C-terminal tail relocates to form contacts with ArpC4 and Arp2, providing further stabilization to the active conformation in the branch junction.

In summary, our structure provides a more detailed understanding of the structural transformations and interactions being formed by the Arp2/3 complex upon branch junction formation. Our actin filament Arp2/3 complex branch junction model also poses exciting new questions and can serve as a platform for obtaining an even more thorough understanding of the mechanisms and function of this crucial actin filament nucleator, by guiding mutagenesis and biochemistry experiments. Obtaining structures from lamellipodia of migrating cells offers great potential for combining *in situ* structural biology experiments of regulatory factors of the actin cytoskeleton with their dynamic studies combining cell biology and genetics. Continued developments in cryo-EM hardware and software will allow obtaining even higher resolution structures of the Arp2/3 complex within cellular environments, from both wildtype cells and cells lacking specific Arp2/3 complex subunit isoforms or regulatory proteins.

## Acknowledgments

This research was supported by the Scientific Service Units (SSUs) of IST Austria through resources provided by Scientific Computing (SciComp), the Life Science Facility (LSF), the BioImaging Facility (BIF) and the Electron Microscopy Facility (EMF). We also thank Dimitry Tegunov (MPI for Biophysical Chemistry) for helpful discussions about the M software, and Michael Sixt (IST Austria) and Klemens Rottner (Technical University Braunschweig, HZI Braunschweig) for critical reading of the manuscript. We also thank Gregory Voth (University of Chicago) for providing us the MD-derived branch junction model for comparison.

## Author contributions

Conceptualization: FKMS; Methodology: FF, FKMS; Software: FF, WW; Validation: FF, FKMS; Formal analysis: FF, FKMS; Investigation: FF, GD, VVH; Data curation: FF, FKMS; Writing – original draft: FF, FKMS; Writing – review & editing: FF, GD, VVH, FKMS; Visualization: FF, FKMS; Supervision: FKMS; Project administration: FKMS; Funding acquisition: FKMS.

## Data and materials availability

The EM-density map of the actin-filament Arp2/3 complex branch junction and one representative tomogram have been deposited in the EMDB (https://www.ebi.ac.uk/pdbe/emdb/) under accession numbers EMD-XXXXX and EMD-YYYYY, respectively. The model of the actin filament Arp2/3 complex branch junction was deposited in the PDB (https://www.rcsb.org/) under accession code PDB-ZZZZ.

## Materials and Methods

### Cell culture

Wildtype Mus musculus NIH-3T3 (RRID:CVCL_0594) fibroblast cells (kindly provided by Michael Sixt, IST Austria) were cultured in Dulbecco’s modified Eagle’s medium (DMEM GlutaMAX, ThermoFischer Scientific, #31966047), supplemented with 10% (v/v) fetal bovine serum (ThermoFischer Scientific, #10270106) and 1% (v/v) penicillinstreptomycin (ThermoFischer Scientific, #15070063). Cells were incubated at 37°C and 5% CO_2_.

Prior to Rac1Q61L transfection (plasmid kindly provided by Vic Small [20]) using Lipofectamine LTX with Plus Reagent (ThermoFischer Scientific, #15338030), NIH-3T3 cells were seeded at 75% confluency in a 6-well plate and incubated at 37°C and 5% CO_2_ for 4h. The primary transfection mix consisting of 2μg of plasmid DNA encoding for Rac1Q61L, 200μl of DMEM and 2μl of PlusReagent was incubated at room temperature (RT) for 10min. 2μl of LTX reagent was added and the mix was incubated at for another 30min. The transfection mix was added dropwise to the cells. Cells were incubated at 37°C and 5% CO_2_ for 16h prior to trypsinization and seeding onto 200 mesh gold holey carbon grids (R2/2-2C; Quantifoil Micro Tools), which were placed in 3D printed grid holders [39]. Prior to seeding, grids were glow discharged in an ELMO glow discharge unit (Cordouan Technologies) for 2 minutes and subsequently coated with 50μg/mL fibronectin (Sigma-Aldrich, #11051407001) in PBS for 1 hour at RT. Cells were allowed to settle on grids for 4h at 37°C and 5% CO_2_ before the culture medium was exchanged with three washing steps with DMEM. Starvation was conducted at 37°C and 5% CO_2_ for 3h before the cells were extracted and fixed as previously described [20]. In brief, grids were placed in a drop of cytoskeleton buffer (10mM MES, 150mM NaCl, 5mM EGTA, 5mM glucose and 5mM MgCl2, pH6.2) with 0.75% Triton X-100 (Sigma-Aldrich, #T8787), 0.25% glutaraldehyde (Electron Microscopy Services, #E16220) and 0.1μg/mL phalloidin (Sigma-Aldrich, #P2141) and incubated for 1 minute at RT. Grids were post-fixed in a drop of cytoskeleton buffer containing 2% glutaraldehyde and 1μg/mL phalloidin for 15 minutes at RT, before being subjected to vitrification.

### Cryo-electron microscopy

Samples were vitrified using a Leica GP2 plunger (Leica Microsystems) set to 4°C and 80% humidity. After transfer into the blotting chamber, excess fixation solution was manually blotted off and 3μl of a solution of 10nm colloidal gold coated with BSA in PBS was added onto the grids. The grids were then vitrified in liquid ethane (−185°C) after backside blotting (3s) using the blotting sensor of the Leica GP2. Samples were stored under liquid nitrogen conditions until imaging.

Cryo-electron tomograms were acquired on a Thermo Scientific Titan Krios G3i TEM equipped with a BioQuantum post column energy filter and a K3 camera (Gatan) using the SerialEM package [40]. Low-and medium-magnification montages were acquired for search purposes and for defining areas of interest for subsequent high-resolution tomography data acquisition, respectively. Gain references were collected prior to data acquisition. Microscope and filter tuning were performed using SerialEM and DigitalMicrograph (Gatan) software, respectively. The slit width of the filter was set to 20eV. Tilt series were acquired with a dose-symmetric tilt scheme [41] ranging from −60º to 60º with a 2º increment and a nominal defocus ranging from −1.75 to −5.5μm. The nominal magnification was 42,000x, resulting in a pixel size of 2.137Å. Individual tilt images were acquired as 11520×8184 super resolution movies of seven frames. The calculated cumulative dose was 170e/Å^2^. Data was acquired over three acquisition sessions contributing 38, 12 and 81 tilt-series, respectively, for a total of 131 tilt-series, keeping above described acquisition parameters constant.

### Image processing

Super-resolution movies were aligned on-the-fly during data acquisition using the SerialEMCCD frame alignment plugin. Tilt series were automatically saved as 2x-binned (2.137Å/px) mrc stacks. These stacks were used for image processing during template matching and PCA-based classification. CTFFIND4 [42] was used to perform CTF-estimation on each tilt individually. Images were low-pass filtered according to their cumulative electron dose. The appropriate filters were calculated using an exposure-dependent amplitude attenuation function and published critical exposure constants [43]. Prior to further processing, poor quality tilt images caused for example by grid bars blocking the beam at high tilt-angles were removed. For preprocessing of tilt series (tilt stack sorting, removal of bad tilts, exposure filtering) the tomoman software package was used.

Tilt series alignment of the exposure-filtered tilt images was done using the IMOD software package [44]. Reconstruction of CTF-corrected tomograms was performed in NovaCTF [45]. CTF correction via phase-flipping used a Z-slab thickness of 15 nm. The resulting tomograms were consecutively binned 2x, 4x, and 8x via Fourier cropping.

Initial processing steps including template matching, principal component analysis (PCA)-based classification and subtomogram averaging were performed using the Dynamo package [46] up to the generation of the reference for subtomogram averaging in RELION (Fig. S2).

To generate a starting reference for template matching 1549 branch junctions were manually picked from 37 8x-binned tomograms (17.096Å/px) using the 3DMOD functionality of the IMOD software package. No angles were assigned to the manual picked positions. Cubic subvolumes with an approximate side-length of 700Å were extracted from 2x-binned tomograms and aligned against a structure of the branch junction derived previously from negative stain tomograms [20]. Alignment was done over 5 rounds, using the internal binning in Dynamo to resample subvolumes to 8.548Å/px.

The resulting average was then bandpass-filtered (100-40Å) and used for template matching branch junctions in the entire dataset consisting of 131 tomograms. For cross-correlation calculation during template matching a mask consisting of two cylinders (both with a 140Å radius) covering the branch, mother filament and the daughter filaments were applied. Full 360° angular scanning during template matching was performed around all 3 axes with a sampling step of 10°.

False-positive cross-correlation peaks (i.e. from areas containing grey value outliers) were removed. Subsequently, the 300 positions with the highest cross-correlation value per tomogram were considered for further processing. For PCA classification in Dynamo the corresponding 39,300 subvolumes were extracted from 8x-binned tomograms, split into 3 groups of equal size and processed in parallel to allow for faster computation. Pairwise cross-correlation calculations were performed for all particles within each group before PCA was conducted. 10 eigenvolumes were calculated and a subset of them was chosen to be employed for separating the particles into 10 classes. Class averages containing only actin filaments were discarded, and class averages exhibiting equally strong densities for the whole branch junction, the mother and daughter filament were included for further processing. To this end, the remaining particles from the 3 classification groups were merged again, resulting in a total number of 17,302 subvolumes. The subvolumes were subjected to one round of bin 8 alignment (Fig. S2) and averaging in Dynamo to provide a reference for classification and subtomogram averaging in RELION [47].

The following processing steps were performed in Warp 1.0.7 [48], M 1.0.9 [18] and RELION 3.08 [49,50].

In Warp super resolution frames were binned (resulting pixel size was 2.137Å) for frame alignment and defocus estimation of individual tilts. For tilt-series alignment, the same tilts and alignment parameters as determined in IMOD were employed. Defocus parameters were then again determined for whole tilt-series. Coordinates of the particles determined from the Dynamo PCA calculation were employed for the extraction of 17,146 subvolumes into cubic subvolumes of 240 voxels at a pixel spacing of 2.137Å and the corresponding 3D CTF/wedge models, which also consider radiation damage by accumulated electron dose.

One round of RELION 3D classification was performed, resulting in 14,296 subvolumes. This dataset was then subjected to RELION 3D auto-refine using the average determined in Dynamo (low-pass filtered to 40Å), resulting in a resolution of 11.9Å at the 0.143 criterion. Subsequently multiparticle refinement was performed in M. Tilt series were refined using image warp with a 9×6 grid and volume warp with a 4×6×2×10 grid. Particle poses and angles were refined for one temporal sampling point. These settings were kept for all following refinements in M. Subvolumes were re-extracted from the refined tilt-series and aligned with RELION 3D auto-refine using the result of the previous iteration filtered to 40Å as reference. This cycling between Warp, RELION and M was performed for a total of three rounds ending on the final iteration in M. RELION post-processing was applied to the resulting half-maps yielding a final structure at 9.0Å resolution at the 0.143 FSC criterion, which was sharpened using an empirically determined B-factor of −50 (Fig. S2). Masks employed for resolution estimation encompassed the Arp2/3 complex and all actin monomers contacting it. Since the actin filaments extended to the box edge, we first masked in Dynamo around the region of interest (i.e. the branch junction and surrounding actin monomers). Then RELION was used to generate a mask around this region of interest, low-pass filtering the mask to 15Å, extending it by 5 voxels and applying a soft edge of 10 voxels (Fig. S2).

### Model fitting and data analysis

The crystal structure of the Arp2/3 complex with ATP bound to Arp2 and Arp3 (pdb 1TYQ) [8] and the single particle cryo-em structure of an ATP and Phalloidin bound actin filament (pdb 6T20) [30] were used to generate a model of the actin filament Arp2/3 complex branch junction.

All subunits of the Arp2/3 complex and individual actin monomers (11 in total, 8 within the densities of the mother filament and 3 within the densities of the daughter filament) were placed using the rigid body fitting option in UCSF Chimera [51]. Actin monomers from pdb 6T20 were fitted individually to improve the rigid body fit. Due to the increased flexibility as suggested by the lower resolution in our map for the N-terminal region of ArpC5, rigid body fitting only considered residues 69 to 151 of this subunit.

In most crystal structures of the Arp2/3 complex, subdomains 1 and 2 of Arp2 are not resolved, except for a structure of GMF-bound Arp2/3 complex [5]. In order to generate a complete model of Arp2 for fitting into our electron microscopy density map, we generated a composite Arp2 model consisting of subdomains 1 and 2 from pdb 4JD2 (GMF-bound Arp2/3 complex) and subdomains 3 and 4 from pdb 1TYQ (not having an GMF interaction in the original crystal structure). The exact composition of the resulting model, which was then rigid body-fitted is reported in Table S2.

For ArpC1, the “protrusion” helix formed by the residues 297 to 305 and the surrounding linker region is not present in the 1TYQ model. The helix itself and residues 306-309 were imported from pdb 1K8K [7] and placed in an empty density at the surface of the actin monomer M4. Merging of the two models and adding residues connecting them, was performed in Coot [52]. The source of the primary structure elements of the resulting model is given in (Table S2). Smaller gaps in the models of other individual subunits were bridged by adding the missing residues in Coot and N- or C-terminal regions were trimmed if no fitting density could be found. (Table S2) refers to the implemented changes. Phalloidin models were removed and the 4-methyl-histidines in the actin structure (pdb 6T20) at position 73 were replaced by histidines. The complete assembly containing all Arp2/3 subunits, actin monomers and remaining ligands was merged into a single pdb file. Coot was then used to move residues within clashing areas and to release entanglements between chains. The resulting model was associated to the sharpened map in ChimeraX [53] and hydrogens were added using the addh command. The ISOLDE plugin [24] was employed to restrict the positions of ligands (ATP and Mg^2+^/Ca^2+^ ions from the original models), ArpC5 and the actin monomers M1, M8 and D3. It was further used to manually move parts of the Arp2 domain 2 (residues 37-56), Arp3 C-terminus (residues 405 onward), and to perform a molecular dynamics simulation for the whole model to allow for an initial flexible fitting into the corresponding densities. In a next step ChimeraX was employed to exchange truncated residues present in the original models with their full-length counter parts (Table S2). Using ISOLDE smaller areas were simulated locally to manually alleviate clashes, improve fitting of sub chains and correct for highly improbable geometries. For this, secondary structure restraints were applied whenever necessary to keep α-helices from deteriorating, and positional restraints were applied to keep the actin structures from occupying densities associated with phalloidin. After regional changes were implemented the whole model was activated again via ISOLDE and cooled down to 0K. After all ligands had been removed, the quality of the model was validated using MOLPROBITY [54] as it is integrated in the Phenix Comprehensive validation (Cryo-EM) tool [55] and is given in (Table S3).

Visualization and model analysis were performed with UCSF ChimeraX. For the comparisons of inactive and active conformation shown in figure 2, maps were generated via the molmap command in ChimeraX at described resolution from pdb 1TYQ and our branch junction model, respectively (1TYQ was modified to contain all residues present in our model).

RMSD calculations were performed considering only the C-alpha atoms of the models. Models were aligned and compared in Chimera using the matchmaker and the RMSD command, respectively. PDB2PQR [56] and APBS [57] were employed to calculate electrostatic potential maps, which were used to color surfaces in ChimeraX.

**Fig. S1.:**
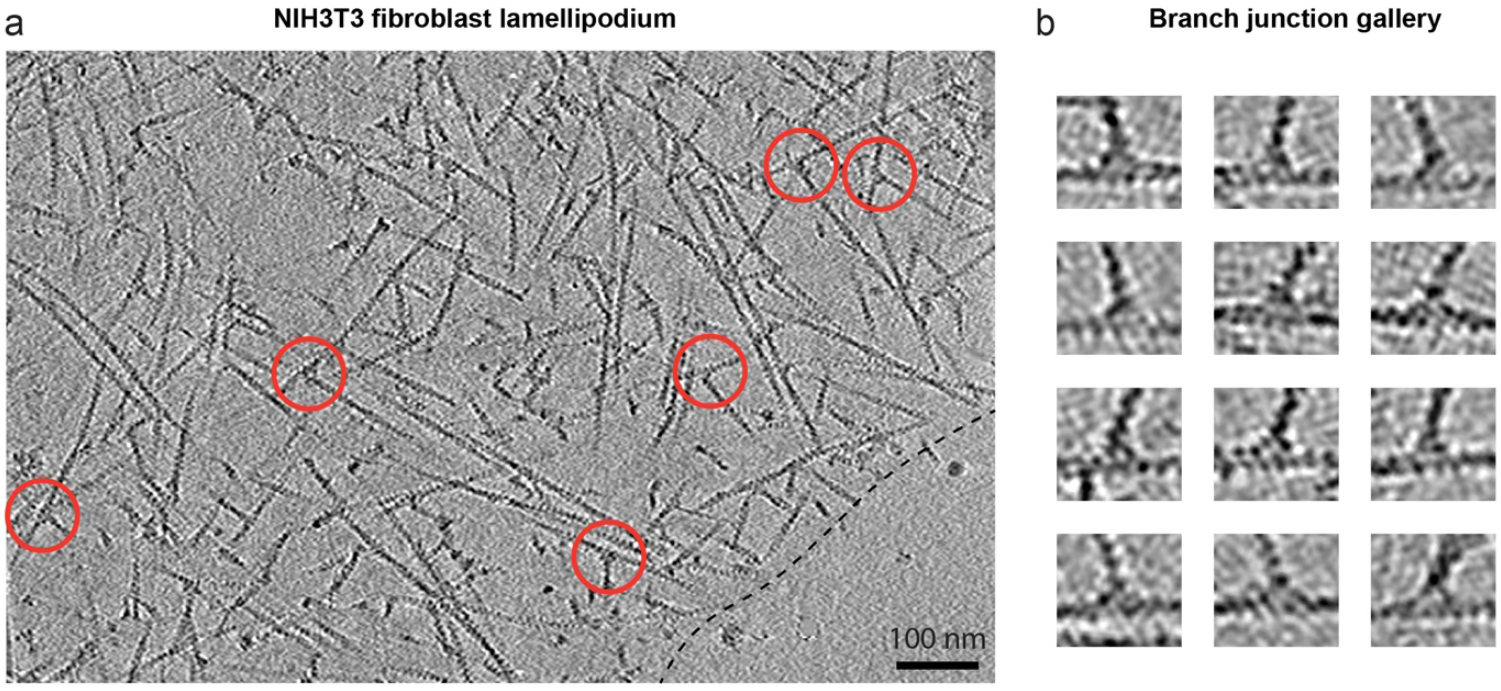
Cryo-ET of the actin network in a NIH3T3 fibroblast lamellipodium. **(a)** Computational slice through a binned, non-CTF corrected tomogram (17.096Å/px) of a NIH3T3 fibroblast lamellipodium. Protein density is black. The dense actin network is visible and the helical appearance of individual actin filaments can be clearly appreciated. Several branch junctions are highlighted in red circles. The dashed line indicates the cell edge. Scale bar is 100nm. **(b)** Gallery of selected branch junctions. The density corresponding to the Arp2/3 complex at branch junctions is visible in the individual examples.

**Fig. S2.:**
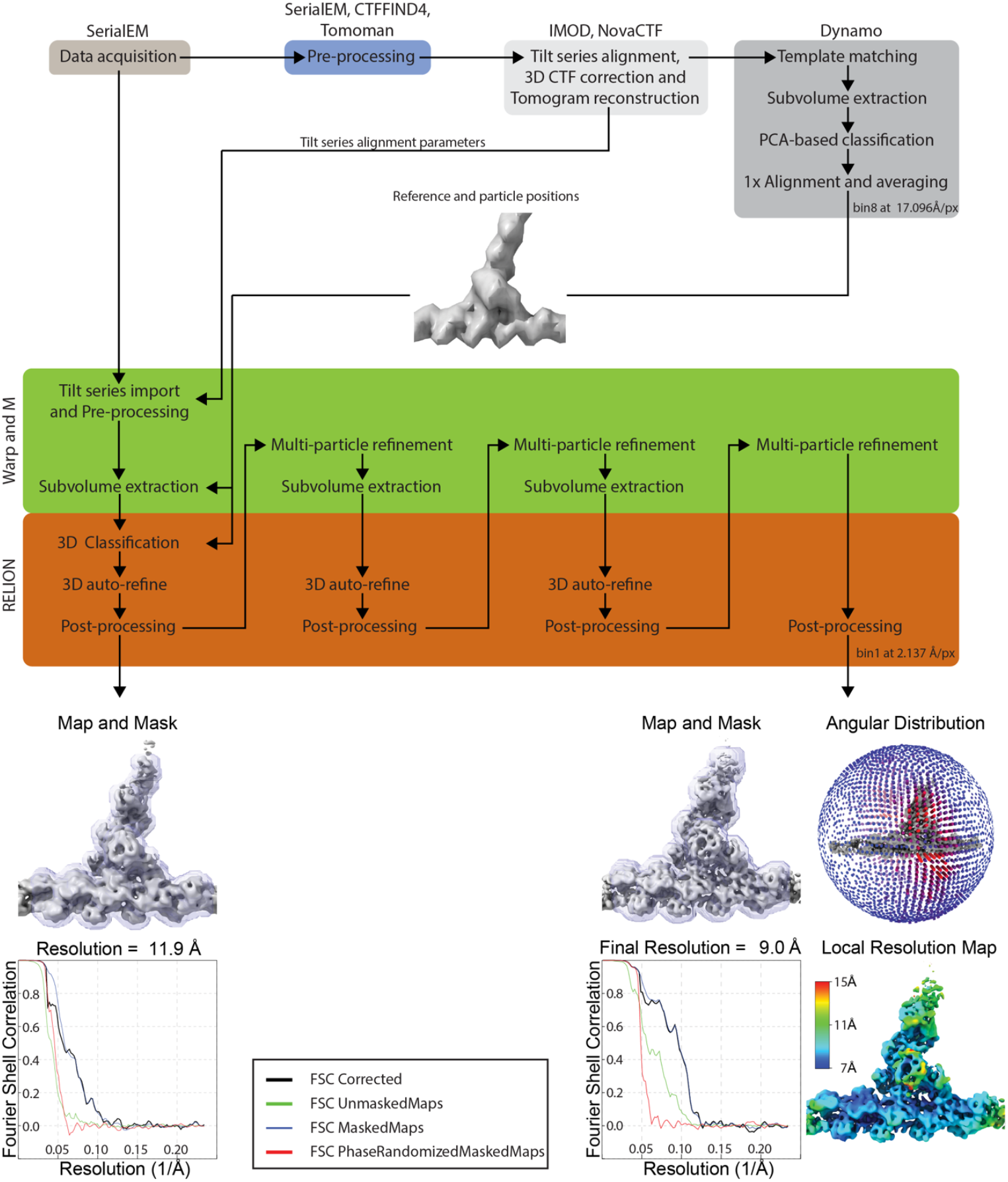
Graphical workflow of image processing. Flow chart indicating the data processing steps involved in generating the structure of the active Arp2/3 complex within the branch junction. Colored boxes indicate usage of specific software packages. For simplicity, the procedure to produce a reference for template matching via manual picking and averaging branch junctions is not depicted here, but is described within the material and methods section.

**Fig. S3.:**
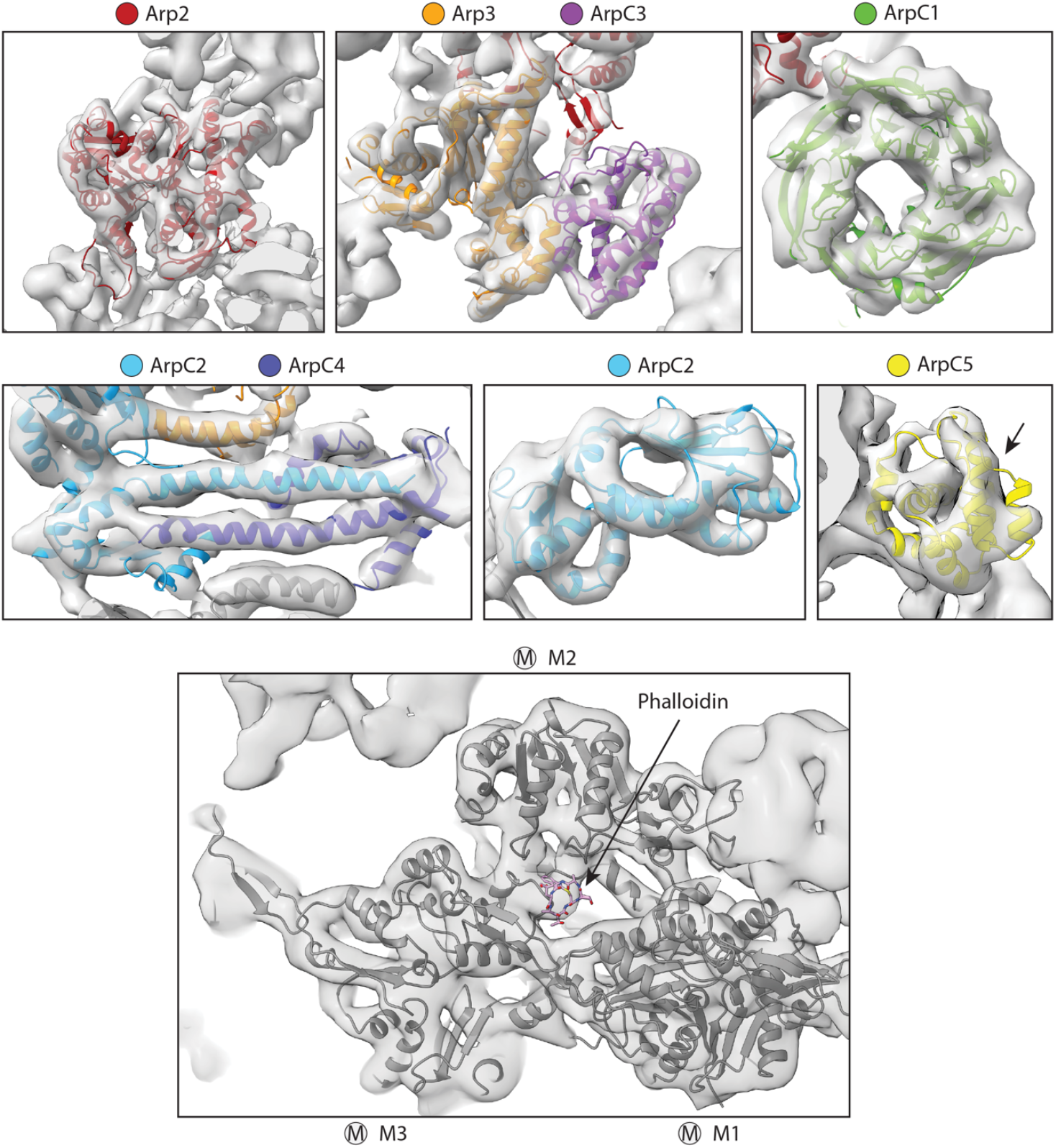
Structural details of the actin filament Arp2/3 complex branch junction. Densities for the individual subunits of the Arp2/3 complex plus their fitted models are shown. Subunit colors are annotated and identical to the schematic guide given in Figure 1. Secondary structure detail (i.e. alpha helices) are clearly visualized, for example for Arp3, ArpC2 and ArpC4. The beta-propellers of ArpC1 allow unambiguous fitting of the subunit into the density of the branch junction. The increased apparent flexibility at the N-terminus of the ArpC5 helical core (annotated by an arrow) is visible, resulting in reduced density for the N-terminal helices of this subunit. The subnanometer resolution of our structure also allows to clearly visualize secondary structure details in the actin filament, further highlighted by the visibility of an additional density accommodating Phalloidin (annotated by an arrow).

**Fig. S4.:**
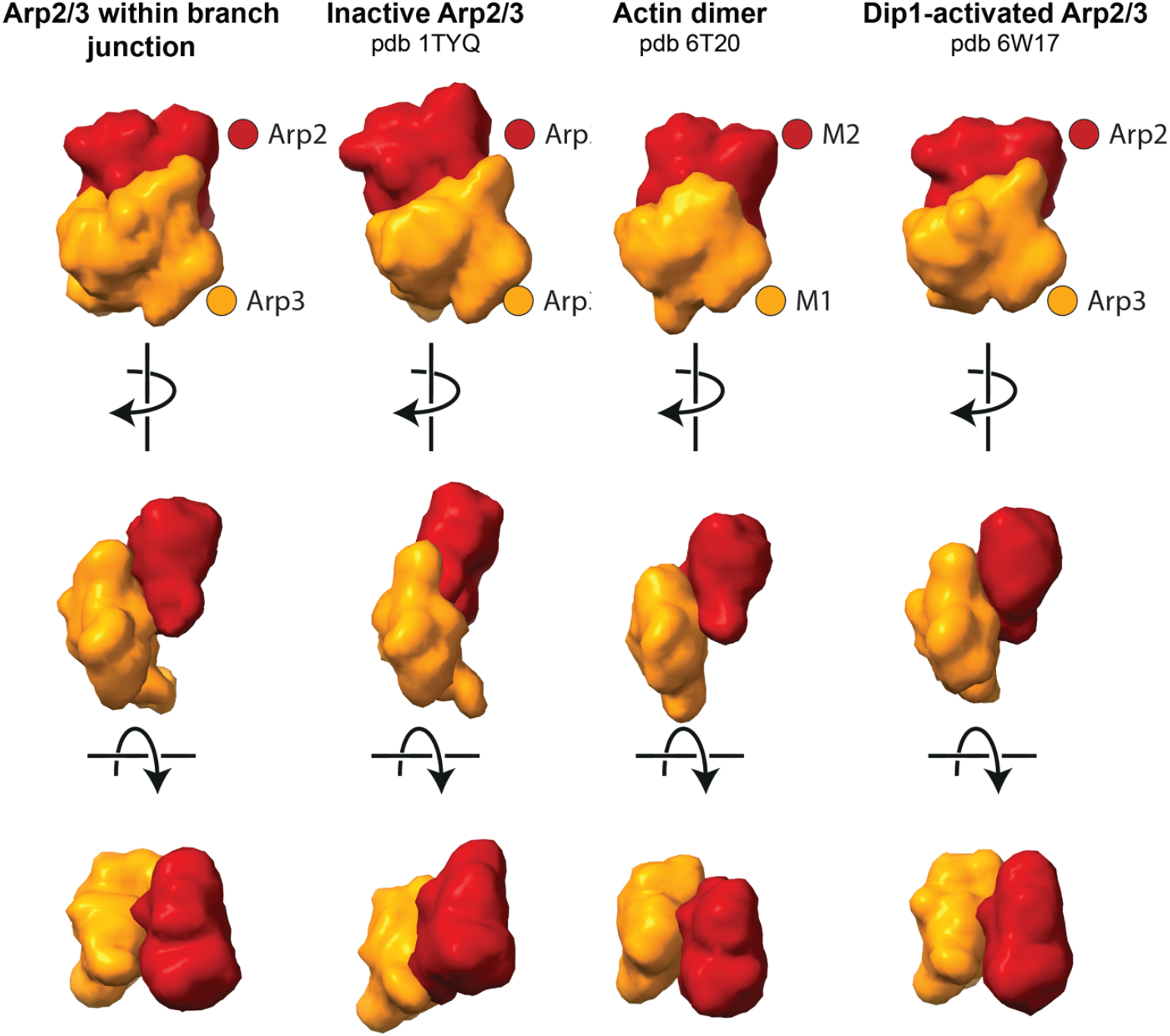
Comparison of the short pitch actin dimer conformation to Arp2 and Arp3 in their inactive and active conformation. Density maps of molecular models of an actin dimer, or of Arp2 and Arp3 in their active and inactive conformation, respectively, filtered to a resolution of 15Å. The models of the active Arp2/3 conformation were derived from our model of the actin filament Arp2/3 complex structure in cells described in this manuscript and the *in vitro* Dip1-activated *S. pombe* Arp2/3 complex (pdb 6W17) [29]. The shown models of filamentous actin were derived from pdb 6T20 [30], and for the inactive ATP-bound Arp2/3 complex from pdb 1TYQ [8]. Maps were oriented by fitting Arp3 subunits and the actin monomer M1 to each other. While Arp2 and Arp3 of the active complex adopt a similar short pitch conformation as the monomers of the actin dimer, this is not the case for the subunits of the inactive complex. Subunit identity is indicated by the color scheme.

**Fig. S5.:**
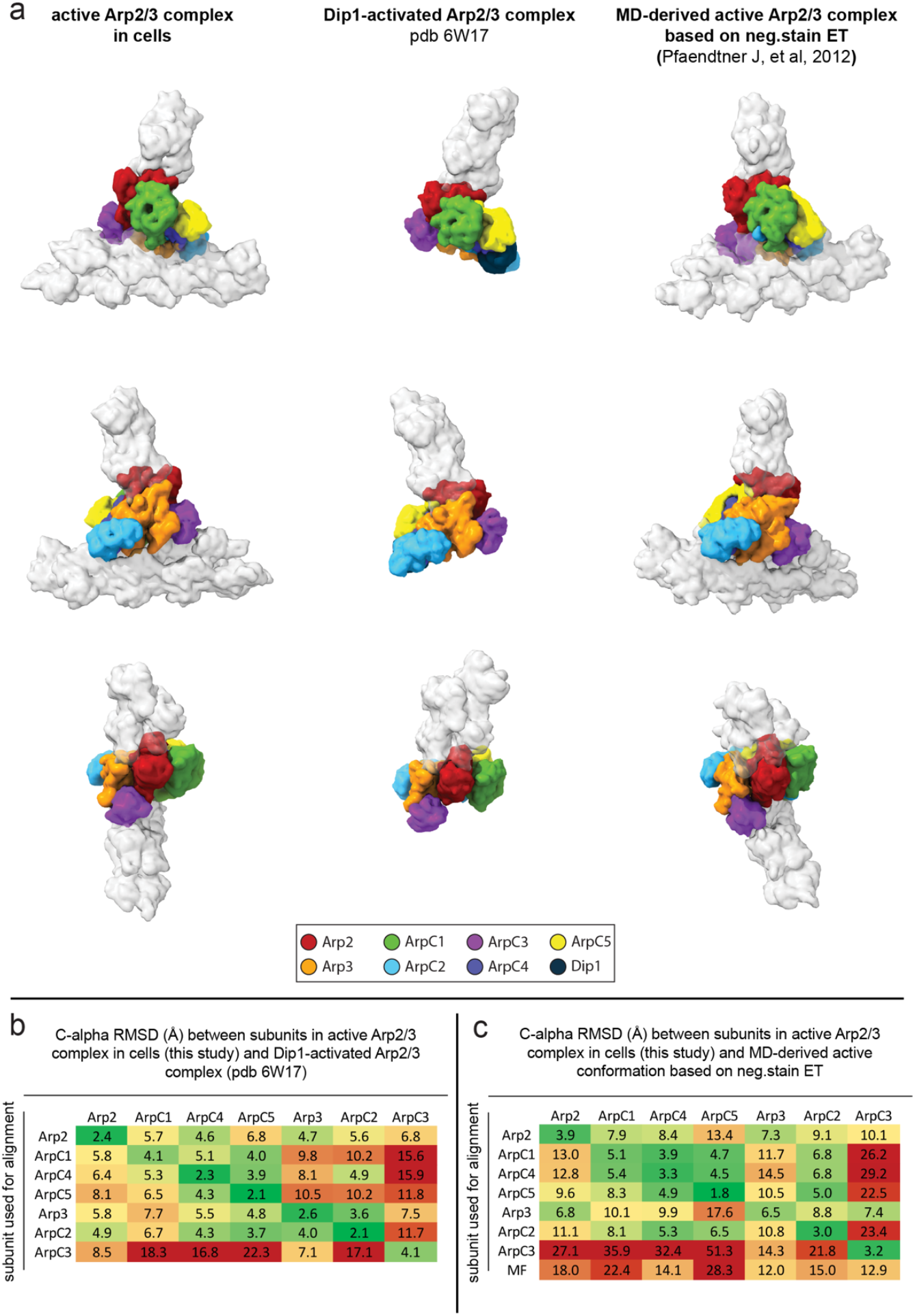
Comparison of the active Arp2/3 complex conformation in branch junctions in cells to the *in vitro* Dip1-activated Arp2/3 complex and to a previously published MD-derived *in vitro* branch model. **(a)** Molecular models of the Arp2/3 complex in the active conformation (shown as density maps filtered to 9.5Å resolution) as observed in cells (left, this study), the *in vitro* Dip1-activated Arp2/3 complex (middle) [29], and a MD-derived active Arp2/3 complex [12], which is based on a low-resolution negative stain ET reconstruction (right) [9]. The models were aligned on the ArpC2 subunits to visualize the different conformations of the Arp2/3 complex and in case for the branch junction also the varying position on the mother filament between the *in situ* and *in vitro* model. **(b-c)** RMSD values (in Å) calculated between the subunits of the three models shown in **(a)**. Rows indicate which subunit was used for aligning full models to each other, prior to measurements between individual subunits (indicated in the columns) of the different models of the active Arp2/3 complex. **(b)** RMSD values of differences between the Arp2/3 complex in branch junctions in cells and the *in vitro* Dip1-activated Arp2/3 complex. In order to calculate C-alpha RMSD values between our model and pdb 6W17 only primary structure areas are considered, in which the *B. Taurus* (as used in our model) and *S. pombe* protein sequences are in the same register, hence omitting inserts present in only one species. This comparison reveals differences between the Dip1-activated Arp2/3 complex in vitro and the activated state of the Arp2/3 complex in cells, in particular with respect to ArpC3. **(c)** RMSD values of differences between the Arp2/3 complex in branch junctions in cells and the MD-derived *in vitro* branch junction model.

**Fig. S6.:**
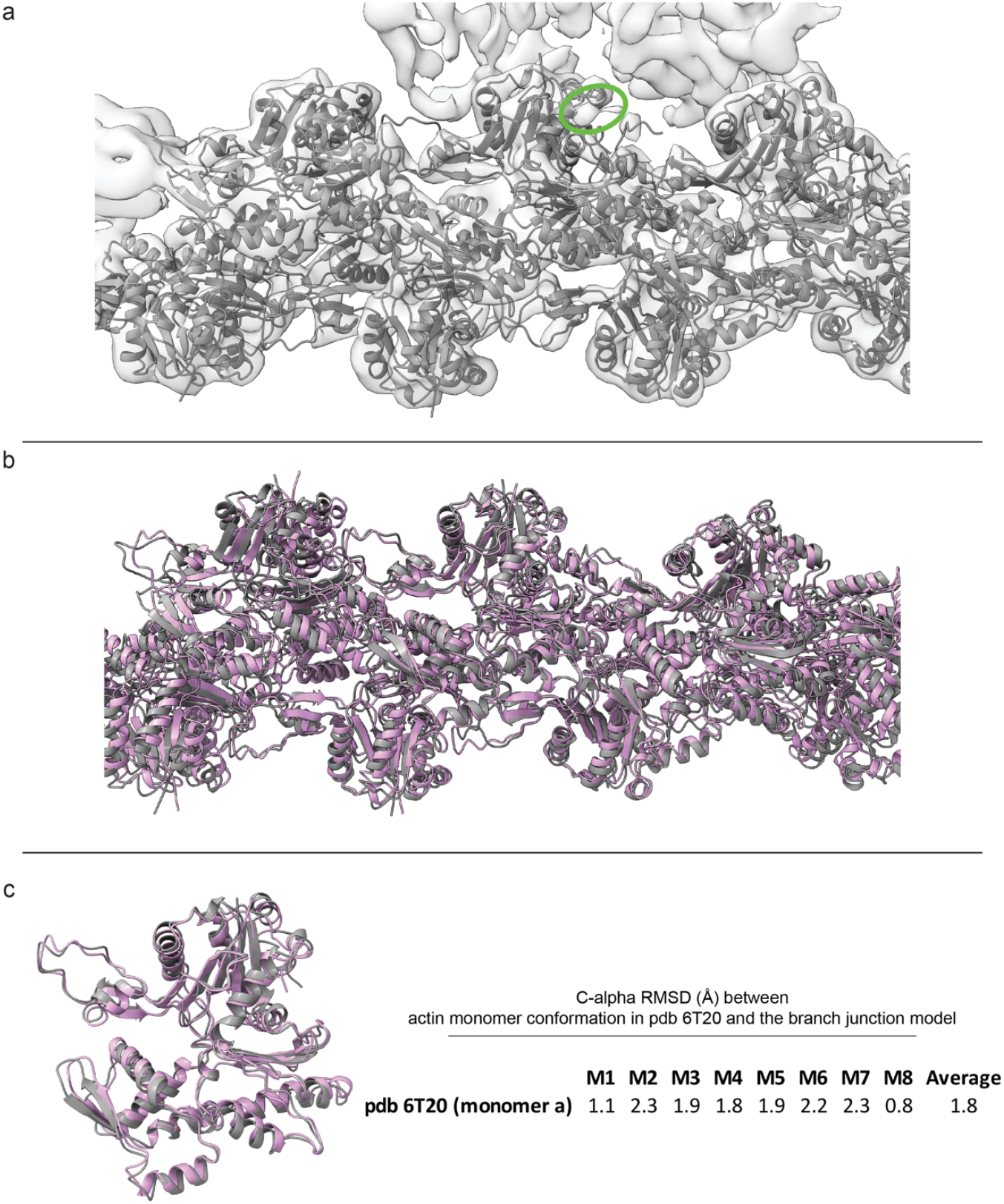
Mother actin filament conformation in the actin-Arp2/3 complex branch junction. **(a)** Fit of the final model of the mother actin filament after MD-refinement into the EM density of the branch junction using ISOLDE. The empty density close to M4, corresponding to the helix of ArpC1 is annotated with a green ellipsoid. **(b)** Superimposition between the starting model used for fitting (pdb 6T20, pink) and the final model after MD-refinement. Small deviations between the filament assemblies can be observed. **(c)** Superimposition between one monomer of pdb 6T20 (pink) and monomer M4 of the mother filament in our branch junction model. No large-scale deviations in the monomer conformation are observed. **(d)** RMSD calculations between the C-alpha atoms of one monomer of pdb 6T20 and the monomers of the mother filament. The average RMSD value is 1.76Å.

**Table S1.** Data acquisition and image processing parameters

| Sample | NIH3T3 fibroblast lamellipodia |  |
| --- | --- | --- |
| Acquisition settings | Microscope | FEI Titan Krios |
|  | Voltage (keV) | 300 |
|  | Detector | Gatan BioQuantum K3 |
|  | Energy-filter | Yes |
|  | Slit width (eV) | 20 |
|  | Super-resolution mode | Yes |
|  | Å/pixel | 2.137 |
|  | Defocus range (microns) | -1.75 to -5.5 |
|  | Defocus step (microns) | 0.25 to 0.5 |
|  | Acquisition scheme | -60/60°, 2°, dose symmetric |
|  | Total Dose (electrons/Å<sup>2</sup>) | ~170 |
|  | Dose rate (electrons/pix/sec) | 19.25 |
|  | Frame number | 7 |
|  | Tomogram number | 131 |
| Processing settings | Subvolumes after template matching | 39,300 |
|  | Subvolumes final | 14,296 |
|  | Final resolution (0.143 FSC) in Å | **9** |

**Table S2.** Summary of model content. The pdb files used to generate the final model of the active Arp2/3 complex are listed. Residues that had to be added or removed from the original models are indicated. Residue stubs in the original models were completed to contain their entire side chains for MD-modelling in ISOLDE.

| Subunit | pdb 1TYQ | residues added with coot | residues added from other source (source) | residues cropped compared to 1tyq | residue stubs completed for side chains for modeling in ISOLDE |
| --- | --- | --- | --- | --- | --- |
| Arp2 | 151-345 | 1, 2, 3 | 4-150, 346-387 (4JD2) |  | 27, 36, 42, 53, 106, 118, 125, 153, 173, 174, 175, 180, 181, 290, 291, 292, 294, 295, 296, 329, 330, 331, 336, 339, 340, 341, 345, 366, 380, 383, 384 |
| Arp3 | 2-39, 52-353, 360-418 | 1, 40-51, 354-359 |  |  | 79, 262, 263, 264, 265 |
| ArpC1 | 1-288, 319-372 | 289-296, 310-318 | 297-309 (1k8k) |  | 1, 130, 131 |
| ArpC2 | 1-208, 217-281 | 209-216 |  | 282 | 207 |
| ArpC3 | 2-150, 155-174 | 1, 151-154, 175-178 |  |  |  |
| ArpC4 | 3-168 | 1, 2 |  |  |  |
| ArpC5 | 36-151 |  |  | 10-27, 35 | 74 |

**Table S3.** Modelling parameters and statistics

| **ISOLDE Refinement** | **Actin filament Arp2/3 complex**<br>**branch junction** |
| --- | --- |
| **Resolution of map (Å)** | 9 |
| **Number of Arp2/3 complex subunits/**<br>**actin monomers** | 7/11 |
| **All-atom clash score** | 3.13 |
| **Rotamer outliers** | 6.7% |
| **Ramachandran outliers** | 2.51% |
| **Ramachandran favoured** | 84.21% |
| **Rmsd (angles, degrees)** | 2.209 |
| **Rmsd (bonds, Å)** | 0.013 |

**Table S4.**
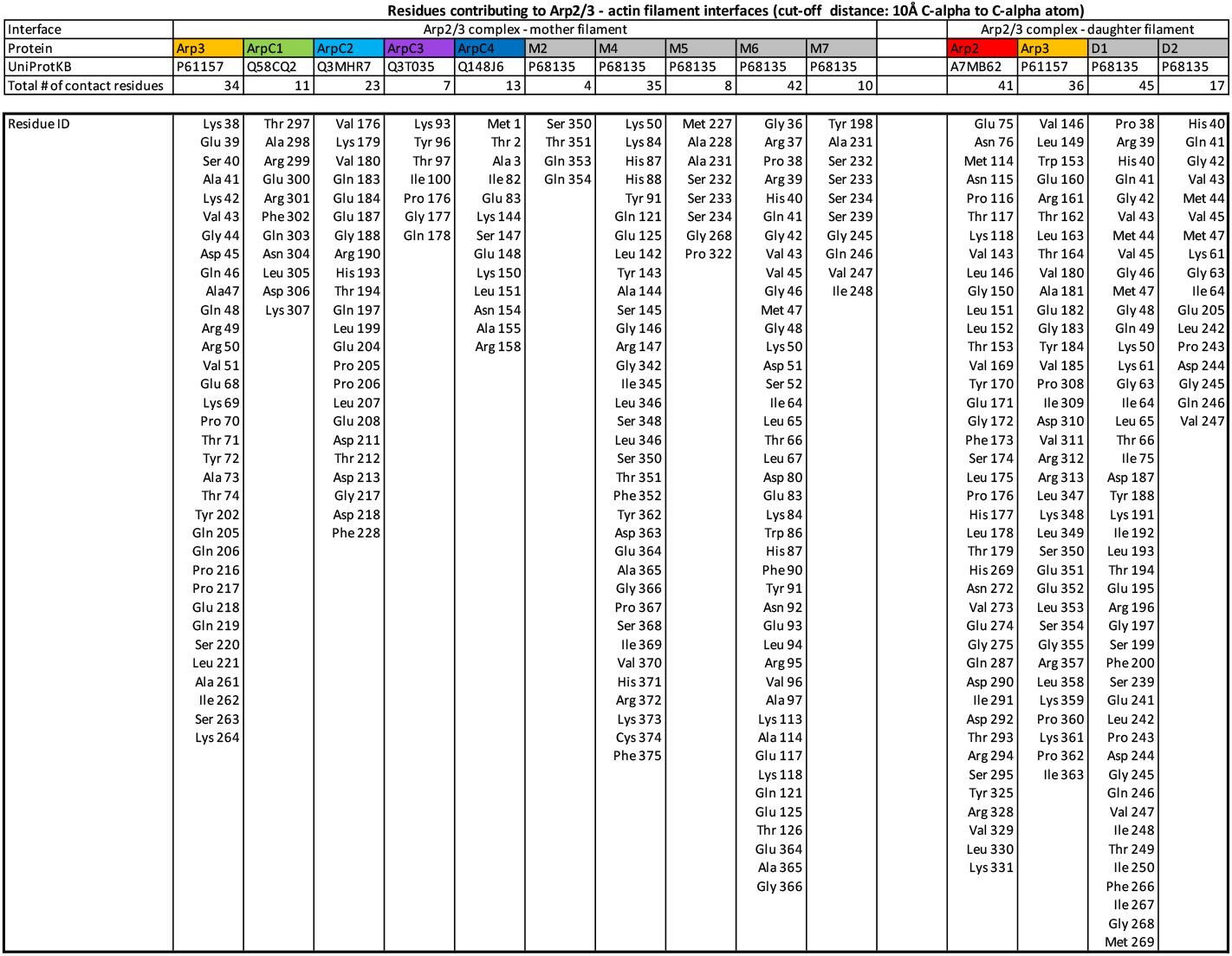
Arp2/3 complex residues contacting the actin mother and daughter filaments. Summary of residues forming interactions between the Arp2/3 complex and the actin filaments, defined by a 10Å C-alpha to C-alpha distance cutoff. The UniProt identifiers for the individual proteins are given.

**Movie S1.**

Tomogram sequence showing the lamellipodial actin network in NIH3T3 fibroblasts

The area shown in this movie corresponds to the actin network shown in Fig. S1.

**Movie S2.**

Rotating view of the actin filament Arp2/3 complex branch junction EM density

**Movie S3.**

Structural tour through the actin filament Arp2/3 complex branch junction

**Movie S4.**

Transition between inactive and active state, barbed end view

The movie shows the transition of the Arp2/3 complex from the inactive state (pdb 1TYQ) to its active conformation. The structure is shown from the barbed end view of the daughter filament.

**Movie S5.**

Transition between inactive and active state, pointed end view

The movie shows the transition of the Arp2/3 complex from the inactive state (pdb 1TYQ) to its active conformation. The structure is shown from the pointed end view of the daughter filament.

